# HLA-E restricted, HIV-1 suppressing, Gag specific CD8+ T cells offer universal vaccine opportunities

**DOI:** 10.1101/2020.12.01.406066

**Authors:** Hongbing Yang, Margarida Rei, Simon Brackenridge, Elena Brenna, Hong Sun, Shaheed Abdulhaqq, Michael K P Liu, Weiwei Ma, Prathiba Kurupati, Xiaoning Xu, Vincenzo Cerundolo, Edward Jenkins, Simon J. Davis, Jonah B. Sacha, Klaus Früh, Louis J. Picker, Persephone Borrow, Geraldine Gillespie, Andrew J. McMichael

## Abstract

Human leukocyte antigen-E (HLA-E) normally presents a HLA class Ia signal peptide to the NKG2A/C-CD94 regulatory receptors on natural killer (NK) cells and T cell subsets. Rhesus macaques immunized with a cytomegalovirus vectored simian immunodeficiency virus (SIV) vaccine, generated Mamu-E (HLA-E homolog) restricted T cell responses that mediated post-challenge SIV replication arrest in >50% of animals. However, human immunodeficiency virus type 1 (HIV-1) specific HLA-E restricted T cells have not been observed in HIV-1-infected individuals. Here we primed HLA-E restricted HIV-1 specific CD8+ T cells *in vitro.* These T cell clones, and allogeneic CD8+ T cells transduced with their T cell receptors, suppressed HIV-1 replication in CD4+ T cells *in vitro*. Vaccine induction of efficacious HLA-E restricted HIV-1 specific T cells should therefore be possible.

**One Sentence Summary:** CD8^+^ T cells that recognize a Gag peptide presented by HLA-E suppress HIV-1 replication *in vitro*.

---

The non-classical human HLA class I molecule HLA-E regulates NK cells and a subset of CD8+ T cells by presenting a nonamer peptide, residues 3-11 of the classical HLA class Ia signal sequence (typically VMAPRTLVL, VL9), to the inhibitory receptor NKG2A-CD94 and its activating counterpart NKG2C-CD94 *(1, 2).* Because the inhibitory receptor is dominant, NKG2A prevents NK cell mediated lysis of cells that co-express HLA class Ia and HLA-E molecules. Similar major histocompatibility complex (MHC) Ia signal peptides bind to homologous non-classical MHC Ib molecules, H-2Qa-1 in mice and Mamu-E in rhesus macaques (RMs) with the same function. Furthermore, human and rhesus cytomegaloviruses down-regulate expression of classical MHC-Ia molecules to evade CD8+ T cell responses but encode their own VL9 signal peptides to ensure that NK cell attack is blocked *(3).*Regulation of NK cell activity is probably the principal function of the HLA-E and its homologs across species.

MHC-E restricted CD8 T cells are rarely found in many infections. However, HLA-E restricted T cells specific for mycobacterial peptides are relatively abundant in humans *(4–6)* and Mamu-E restricted T cell responses were prominent in RMs immunized with rhesus cytomegalovirus (RhCMV) strain 68-1 vectored vaccines *(7).*Atypical MHC class II restricted CD8+T cell responses were also detected in these vaccinated macaques, while classical class Ia restricted CD8 T cell responses were absent *(8).* When these immunized RMs were challenged a year later with pathogenic SIVmac239, all animals were initially SIV infected, but then more than half stringently controlled and subsequently cleared the SIV infection *(9–11).* Recent studies *(12,13)* show that the Mamu-II restricted T cell responses could not mediate protection, indicating that the MHC-E response stimulated by this vaccine is required for protection. This raises the possibility that HLA-E restricted HIV-1 specific responses might be able to control and subsequently clear HIV-1 early after infection in humans. Although one Gag-derived HIV-1 peptide that could bind HLA-E inhibited NK cells through interaction with NKG2A-CD94 *(14)*, HLA-E restricted CD8+ T cell responses have not been described in people.

All RMs immunized with the RhCMV 68-1 vaccine elicit responses to the SIV Gag peptide RMYNPTNIL (RL9SIV), and this peptide has a close homolog in HIV-1 Gag RMYSPTSIL. RL9HIV-1 binds to HLA-E *in vitro* and its crystal structure has been determined *(15).* Therefore, we chose this peptide as our immunogen and prepared HLA-E RL9 tetramers for detecting and sorting responding T cells. Because there is some instability in this HLA-E peptide complex *(15)* with thermal melt analysis showing multiple thermal transition profiles (Figure S1A), we replaced the bulky tyrosine at position 84 on the HLA-E alpha-1 helix with a cysteine (HLA-EY84C) and extended the carboxyl end of the peptide by a glycine and cysteine (RL9GC). These changes create a covalent disulfide bond to link the peptide to HLA-E, giving a stable refolded protein with a thermal melt derivative (TmD) value of 52.2°C (Figure S1B). Previous studies have shown that this modification does not change the structure of the murine MHC class I protein H-2K^b^*(16).*

CD8+ T cells in peripheral blood mononuclear cells (PBMCs) from six HIV-1 negative blood donors were stained with disulfide trapped HLA-E RL9 tetramer [RL9HIV-(D) tetramer]. HLA-A2 negative donors were chosen as some peptide binding motifs previously described for HLA-E also overlap with those reported for HLA-A2 *(17).* Initial tetramer staining detected a mean of 0.004% of CD8+ T cells in freshly isolated PBMCs (Figure 1A). To expand epitope-specific T cells, PBMCs were cultured for 9 days with autologous dendritic cells that had been differentiated with a cytokine cocktail of GM-CSF and IL-4. The DC maturation stimuli of TNF-α, IL-1β and prostaglandin E2 were added after 1 day together with the RL9 peptide, IL-7 and IL-15 to prime RL9 specific CD8+ T cells. IL-2 was added on day 6 *(18).* After 9 days, cell expansion was monitored using disulfide trapped RL9HIV-(D) tetramer. As shown in Figure 1A; tetramer binding T cells were clearly detectable, although still rare with a mean of 0.019% of CD8+ T cells. In donor HD1, with the highest tetramer-positive T cell frequencies, T cells were sorted by disulfide trapped RL9HIV-(D) tetramer and plated out at 0.4 cells per well in microtiter wells and cultured with PHA, IL-2 and feeders. After two weeks, 171 clones had grown, of which 40 had >2% of cells stained with the disulfide trapped RL9HIV-(D) tetramer (Figure 1C). All RL9 positive clones were CD94 negative and did not stain with tetramers of HLA-E disulfide trapped to the HLA class I signal peptide VL9 [VL9-(D) tetramer] (Figure 1D). Fifteen clones that stained in the tetramer positive range of 10-40% were chosen for further study. When these cells were fixed in 2% paraformaldehyde immediately after tetramer staining but in the absence of washing, close to 100% staining was observed (Supplementary figure S2), suggesting that the lower values detected for unfixed cloned cells were caused by relatively low affinity tetramer binding to T cell receptors (TCR). These clones were then maintained in culture for approximately 2 months and expanded with irradiated feeders and PHA every 15-18 days.

**Figure 1.**
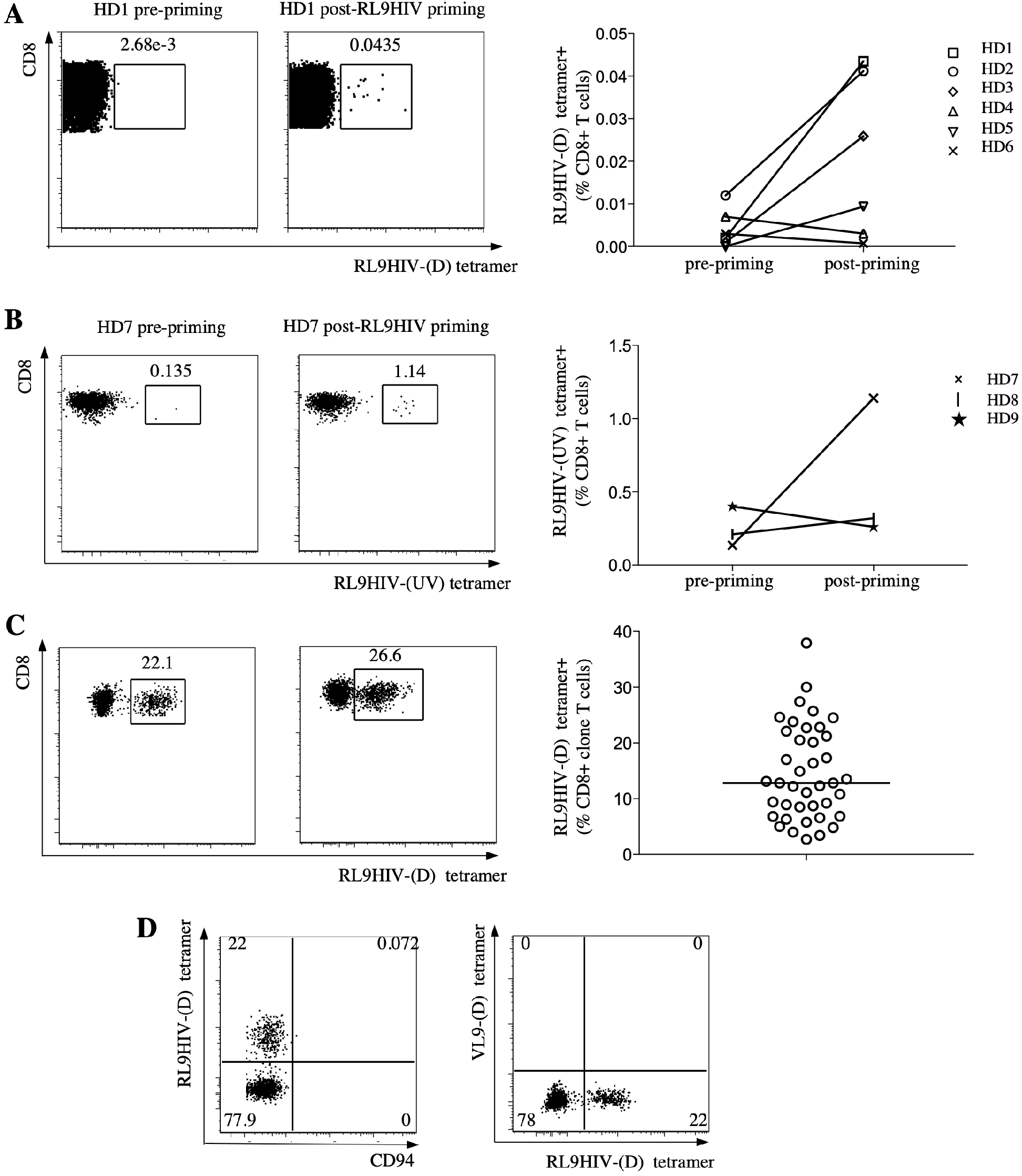
Priming and cloning of HLA-E restricted RL9HIV-specific CD8^+^ T cells from HIV naïve donors. **(A)** PBMCs from 9 HIV-1 seronegative donors (HD 1 to 9) were stimulated with autologous activated dendritic cells and the RL9HIV peptide for 9 days. HLA-E restricted RL9HIV specific CD8+ T cells were identified using HLA-E-RL9 disulfide trapped tetramer (RL9HIV-(D)) for donors HD1 to 6 or (**B**) a HLA-E RL9 tetramer generated by UV exchange RL9HIV (UV)) for donors HD 7 to 9. (**C**) disulfide trapped RL9HIV-(D) tetramer+ cells from donor HD1 were sorted for single cell cloning, and positive clones were identified using disulfide trapped RL9HIV-(D) tetramer. (**D**) RL9 positive clones were CD94 negative and do not recognize HLA-E bound to canonical VL9 signal peptide as illustrated by lack of dual staining with RL9 and VL9 disulfide trapped HLA-E tetramers.

In a separate experiment, to validate the disulfide-trapped HLA-E RL9 tetramer tool, we used an alternative tetramer method involving UV peptide exchange. RL9 specific CD8+ T cells were expanded from PBMC from three additional HIV-1 negative blood donors following the same protocol. For comparison, an HLA-E RL9 nondisulfide trapped tetramer was freshly prepared using an UV-mediated peptide exchange refolding method *(19)*. Here, HLA-E was first refolded stably with the signal VL9 peptide modified to replace position 5 arginine with the light sensitive 3-amino-3-(2-nitrophenyl)-propionic acid residue and then the RL9 peptide was exchanged by exposing this complex to UV light in the presence of excess RL9 peptide *(15)*. Staining of PBMCs with the UV exchanged HLA-E RL9 tetramer [RL9HIV-(UV) tetramer] prior to expansion showed a mean of 0.25% tetramer-positive, a value higher than that observed in the first 6 donors studied due to slightly higher non-specific binding of this tetramer compared to the disulfide trapped RL9HIV-(D) tetramer (Figure 1B). One donor, HD7 showed a marked increase in the non-disulfide trapped tetramer positive population after priming (0.2% to 1.14%) (Figure 1B). These findings show that the presence of HLA-E RL9 specific T cells in our original donor was not unique, and that irrespective of the tetramer reagent used, antigen specific T cells could be enriched from healthy donor PBMCs.

The functional capacities of the 15 clones were tested by mixing with (HLA-negative) K562 cells expressing a disulfide trapped single chain trimer of HLA-E-β2m-RL9 (HLA-D-RL9), which was shown to stimulate degranulation (CD107a/b expression), secretion of TNFα and IFNγ and CD137 expression (Figures 2A, S3). These responses were compared to those elicited by K562 cells expressing HLA-E-β2m-RL9 as a single chain trimer, but non-disulfide trapped (HLA-E-RL9) *(1, 20)* and by K562 cells transduced with just the HLA-E heavy chain, expressing with endogenous ß2m, pulsed with RL9 peptide. Responses to these last stimuli were much weaker than those elicited by cells expressing the disulfide trapped HLA-E-β2m-RL9 single chain trimer (HLA-D-RL9) (Figure 2B). CD107a/b up-regulation was more readily elicited than TNFα production with most clones demonstrating significant responses to RL9 peptide pulsed K562-E cell line and non-disulfide trapped K562-E-RL9 single chain trimer (Figure 2B). Clone, p13c7 gave a measurable response to RL9 peptide pulsed cells and the TNFα and CD107a/b responses generated by this clone were blocked by competitive inhibition with the canonical HLA-E binding VL9 signal peptide, confirming that RL9 was presented by HLA-E (Figure 2C).

**Figure 2.**
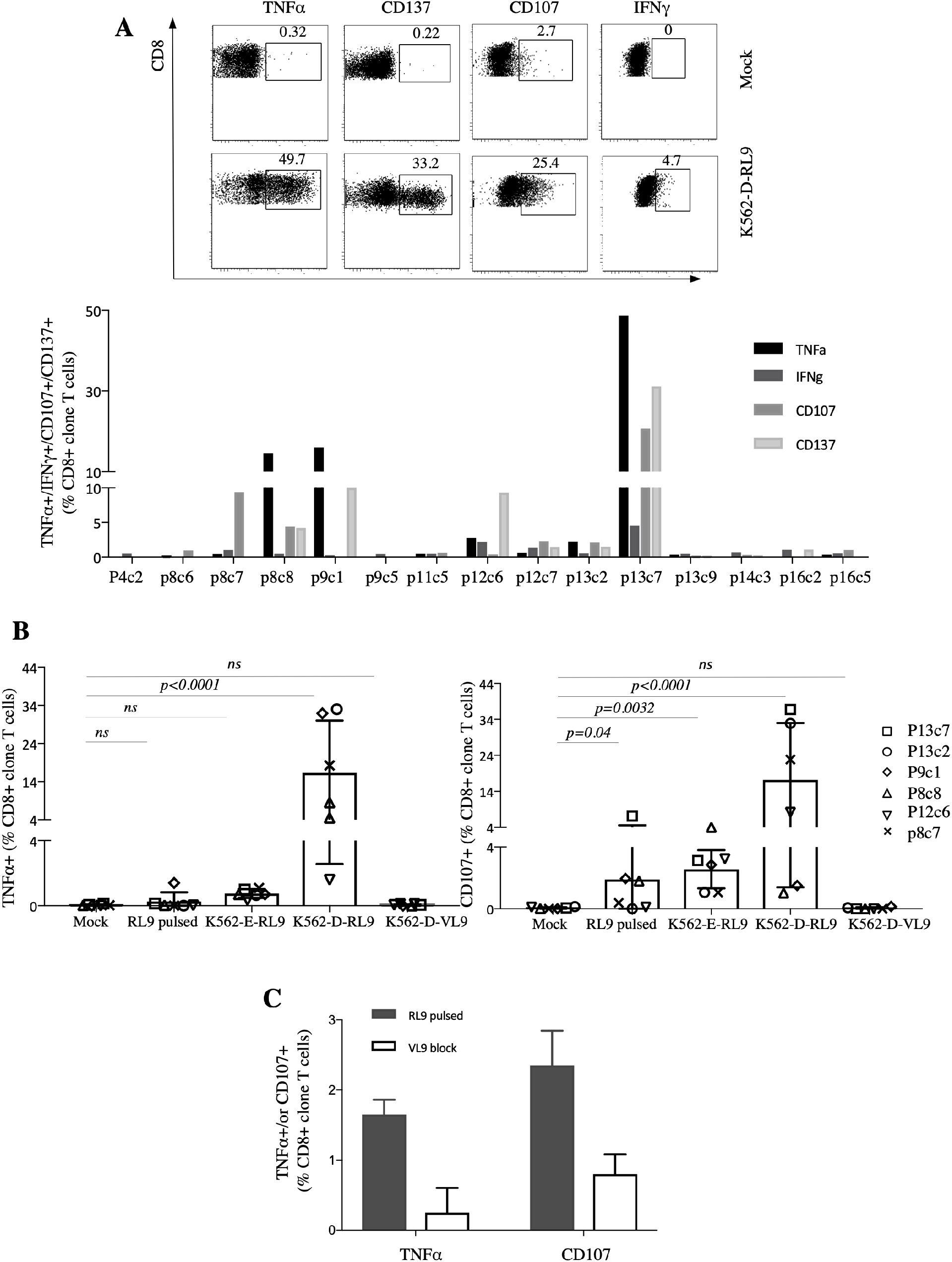
Functional analysis of HLA-E restricted RL9HIV-specific CD8^+^ T cell clones. **(A)**IFN-γ, TNF-α, CD107a/b and CD137 expression by 15 RL9 clones upon stimulation with K562 cells transduced with single chain trimer (SCT) disulfide trapped HLA-D-RL9 were assessed using flow cytometry-based readouts; responsive clones could be detected using multiple functional readouts. (**B**) Six positive clones were further assessed with comparison of stimulation with K562 transduced with HLA-E and pulsed with RL9 peptide, K562 transduced with SCT linked HLA-E RL9 (K562-E-RL9), K562 transduced with SCT disulfide-linked HLA-E RL9 (K562-D-RL9), and K562 transduced with SCT disulfide linked HLA-E VL9 (K562-D-VL9) as a negative control. Horizontal lines indicated means. Groups were analyzed by 2-way ANOVA with Tukey’s multiple comparisons. Data shown are representative of six donors and two independent experiments. (**C**) Blockade of TNF-α secretion and CD107 expression of clone p13c7 by addition of the VL9 peptide prior stimulation with HLA-E transduced K562 cells pulsed with RL9 peptide. Data shown are from two independent experiments.

Given that T cells were primed *in vitro* with the 9mer RL9 peptide and predominantly tested with the disulfide trapped RL9HIV-(D) tetramer, it was important to check whether the clones recognized HIV-1 infected cells. Therefore we set up a viral inhibition assay (VIA), as used previously for the classical MHC Ia restricted CD8 T cells *(21),* where CD4-expressing 721.221 HLA-A and B deficient, HLA-E positive cells were infected with HIV-1 NL4.3 and then incubated with test T cell clones at an E:T ratio of 1:1 for 5 days. HIV-1-infected cells were detected on the basis of Gag p24 expression and the reduction of Gag p24+ cells indicated inhibition of HIV-1 replication and/or lysis of HIV-1 infected cells, mediated by the specific T cell clone. Six clones were tested, and they reduced the p24 positive cells by 15-45% (Figure 3A). Furthermore, exposure to 721.221-CD4 cells infected with HIV-1 NL4.3 stimulated significant greater up-regulation of surface CD137 *(22, 23)* on the T cell clones compared to that elicited by exposure to uninfected cells *(p=0.031).* This activation of CD8+/CD94-T cells was significantly blocked by addition of the VL9 signal peptide (*p=0.031*), indicating that the T cells recognized peptide bound to HLA-E. A control HLA-B*08 Epstein Barr Virus (EBV) specific CD8+ T cell clone did not respond to HIV-1 infected 721.221-CD4 cells (Figure 3B). We also tested the T cell clones on purified autologous CD4+ T cell targets, stimulated with anti-CD3 for 3 days prior HIV-1 NL4.3 infection. Viral inhibition by clones was again observed, with greater suppression obtained at effector (clone) to target cell ratios of 5:1 compared to 1:1 (Figure 3C).

**Figure 3.**
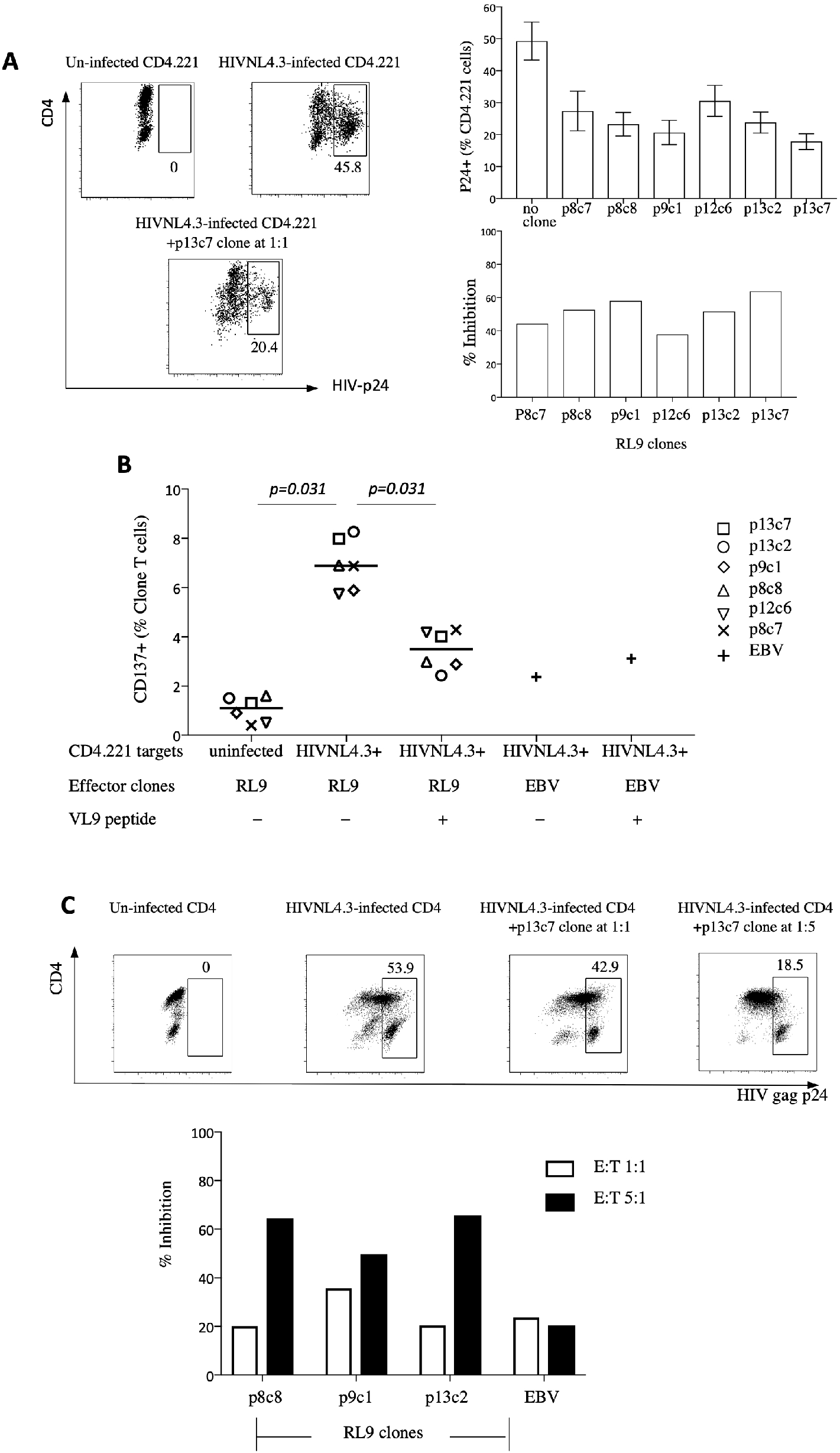
Recognition of naturally presented RL9 epitope on HIV-1 virus infected cells and inhibition of HIV-1-infected targets by RL9 clones. **(A)** 771.221 cell lines transfected with CD4 were infected with HIV-1NL4.3 virus and cultured alone or with RL9-specific CD8+ T cell clones at an E: T ratio of 1:1. Frequencies of HIV-infected cells (Gag p24+) and the percentage reduction in the frequency of Gag p24+ cells in the presence of the RL9 clones after 5 days of culture (calculated as described in Materials and Methods) are shown. (**B**) Significant *(p=0.031)* higher expression of the CD137 activation marker was observed on clone CD8+ T cell clones when co-cultured with HIV-1NL4.3 virus infected CD4.221 compared to uninfected CD4.221 cells, and this was significantly blocked by the VL9 peptide (*p=0.031*). A control EBV-specific CD8+ T cell clone generated from the same donor showed minimal CD137 expression that was not affected by VL9 addition. Horizontal lines indicated means. Data were analyzed by the non-parametric Wilcoxon signed rank test. (**C**) Purified autologous CD4+ T cells were stimulated with anti-CD3 for 3 days prior HIV-1NL4.3 infection, then either cultured alone or with clones at E: T ratios of 1:1 and 5:1 for 5 days. Gag p24+ cells were gated on CD3+/CD8-/CD4+ and CD4-T cells. A reduction in the proportion of HIV-1NL4.3 infected primary CD4+ T cells was observed at an E: T ratio of 5:1.

We sequenced the αβT cell receptors (TCRs) from 13 clones and obtained 12 clone-specific sequence pairs, confirming their clonality (Table 1). The TCR α and βgenes of 5 responsive clones were subcloned, inserted into a lentiviral vector and transduced into a Jurkat T cell line J8, genetically modified to delete their endogenous TCR and to express CD8 rather than CD4 (Figure S4, Table S1); eGFP was expressed under the control of the nuclear factor of activated T cells (NFAT) promoter in these J8 Jurkat cells to provide a readout of T cell activation. The initial TCR transduction gave a low frequency of tetramer positive cells but transduced cells were enriched by subsequent sorting to expand J8 Jurkat sublines where > 20% of cells expressed the TCRs (Figure 4A). TCR-expressing J8 Jurkat cells were then exposed to HLA-E transduced K562 cells pulsed with RL9 peptide, to K562 cells transduced with non-disulfide trapped single chain trimer HLA-E-RL9 and to disulfide trapped single chain trimer K562-D-RL9. Expression of the activation marker CD69 and the reporter eGFP were increased on exposure to RL9 peptide pulsed HLA-E-transduced K562 cells, and to cells expressing both HLA-E RL9 disulfide trapped (HLA-D-RL9) and non-trapped single chain trimers (HLA-E-RL9) (Figure 4B).

**Figure 4.**
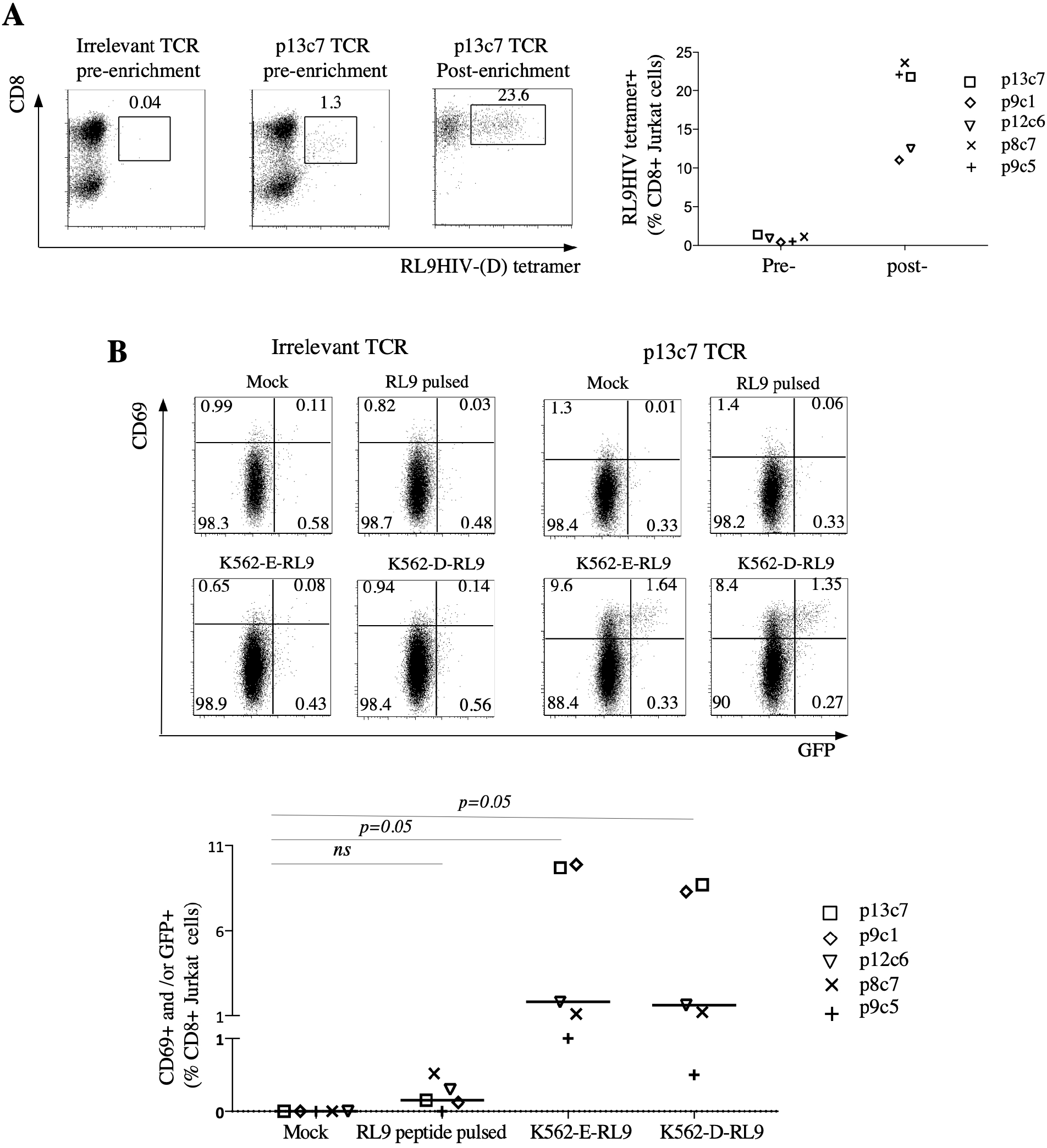
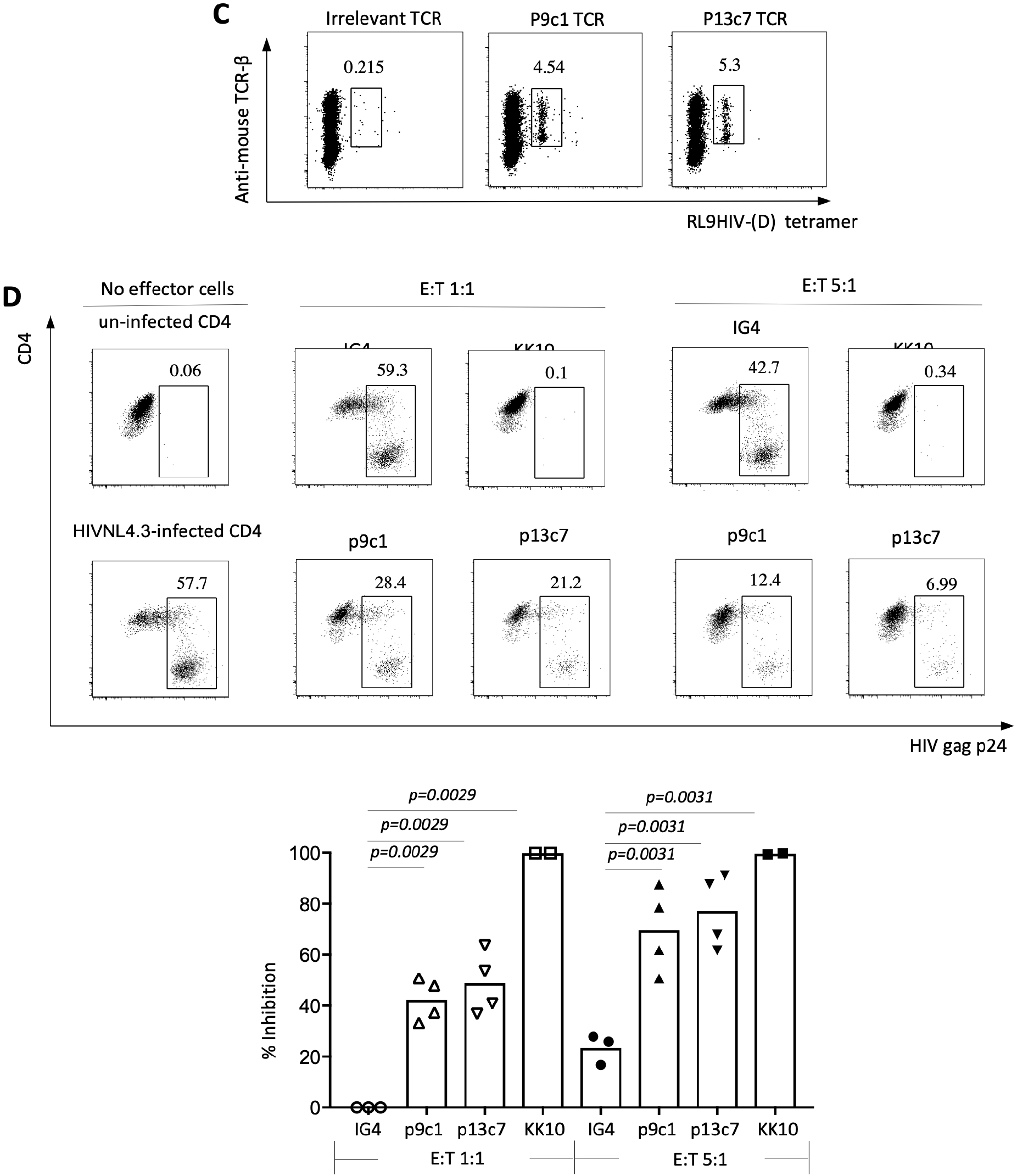
Specificity and function of RL9-specific TCRs transduced into J8 Jurkat and primary CD8+ T cells. **(A)**Five TCR transduced J8 Jurkat cell lines were stained with disulfide trapped RL9HIV-(D) tetramer and the tetramer+ population was enriched by sorting. (**B**) Transduced J8 Jurkat cells (pre-labeled with CellTrace Violet to facilitate gating for subsequent analysis) were partially activated on exposure to HLA-E transduced K562 cells pulsed with RL9 peptide, and more strongly by K562 cells transduced with SCT HLA-E-RL9 or SCT disulfide-linked HLA-D-RL9 as assessed by CD69 and/or GFP up-regulation. Horizontal lines indicated means. Statistical analysis of the data was performed using the nonparametric Wilcoxon signed rank test. (**C**) Two RL9 TCRs p9c1 and p13c7 were transduced into primary CD8+ T cells. CD8+ transductants were stained with disulfide trapped RL9HIV-(D) tetramer initially, washed with PBS, and then stained with anti-mouse TCR Vβ antibody, anti-CD8 and Live/Dead Fixable Aqua. (**D**) A reduction in the percentage of HIV-1NL4.3-infected primary CD4+ T cells was demonstrated when CD8+ transductants were co-cultured with targets at an E:T ratio of 1:1, with greater reduction observed at an E:T ratio of 5:1. Gag p24+ cells were gated on CD3+/CD8-/CD4+ and CD4-T cells. Bars indicate mean reduction in % of p24+ cells. CD8+ T cells transduced with an irrelevant TCR (IG4) were included as a negative control and CD8+ T cells transduced with a TCR recognizing the B*2705 restricted Gag KK10 epitope were included as a positive control when CD4+ T cells from B*2705+ donor were infected with HIV-1NL4.3 virus and used as targets. Statistical analysis of the data was performed by One-way ANOVA Kruskal-Wallis test. Data shown are representative of four independent experiments. The two experiments where KK10 specific TCR transduced CD8+ T cells were tested on HLA B*2705 negative cells and showed no inhibition.

**Table 1.** T cell receptor (TCR) sequences of HLA-E restricted RL9HIV-specific CD8+ T cell clones.

| Clones | CD3 alpha |  | CD3 beta |  |
| --- | --- | --- | --- | --- |
|  | V | J | V | J |
| p4c2 | TRAV38-1 | TRAJ28 | TRBV12-4 | TRBJ1-2 |
| p8c6 | TRAV38-1 | TRAJ43 | TRBV5-6 | TRBJ1-1 |
| p8c7A | TRAV38-1 | TRAJ28 | TRBV9 | TRBJ1-6 |
| p8c7B | TRAV23DV6 | TRAJ57 | TRBV9 | TRBJ1-6 |
| p8c8 | TRAV12-2 | TRAJ48 | TRBV28 | TRBJ2-6 |
| p9c1 | TRAV38-1 | TRAJ45 | TRBV28 | TRBJ2-3 |
| p9c5 | TRAV12-3 | TRAJ29 | TRBV2 | TRBJ1-3 |
| p11c5 | TRAV13-1 | TRAJ16 | TRBV28 | TRBJ2-2 |
| p12c6 | TRAV1-2 | TRAJ34 | TRBV7-9 | TRBJ2-3 |
| p13c2 | TRAV29DV5 | TRAJ22 | TRBV5-1 | TRBJ2-6 |
| p13c7 | TRAV10 | TRAJ17 | TRBV6-6 | TRBJ2-3 |
| p14c3 | TRAV38-2DV8 | TRAJ43 | TRBV19 | TRBJ1-5 |
| p16c5 | TRAV19 | TRAJ40 | TRBV20 | TRBJ2-7 |

Finally, we transduced the TCR Vα/Vß from clones p9c1 and p13c7, fused to murine Cα/Cß, into primary CD8+ T cells. CD8+ T cell transductants were stained with disulfide trapped HLA-E-RL9 tetramers at day 4 post-transfection (Figure 4C) and mouse Cβ-positive cells were sorted to enrich for RL9-specific TCR expressing cells. These were cultured in RPMI 1640 CM (10% AB serum) with IL-15 and IL-2 for a further 17 days before functional analyses. The CD8+ T cell TCR transductants up-regulated CD137 expression and/or produced TNFα when stimulated with either RL9 peptide pulsed autologous EBV transformed B cells (Figure S4A) or HIV-1 NL4.3-infected 721.221-CD4 cells (Figure S4B), and responses were partially blocked by competitive inhibition with the HLA-E binding canonical signal peptide VL9. Then the CD8+ T cell TCR transductants were tested on activated primary CD4+ T cells from healthy allogeneic donors infected with HIV-1 NL4.3 virus, cultured at E: T ratios of 1:1 and 5:1 for 5 days. An irrelevant TCR specific for HLA-A*0201-NY-ESO-1_157-165_ (SLLMWITQC) (IG4) was transduced into CD8 T cells from the same donor and included as a negative control, whilst an HLA B*2705 HIV-1 Gag_263-272_ (KRWIILGLNK) (KK10) specific TCR transduced CD8+ T cells was included as a positive control because the CD4+ T cells were purified from a B*2705+ donor. Both the p9c1 and p13c7 RL9-specific TCR CD8+ transductants significantly diminished HIV-1 virus replication, reducing the %p24+ cells by a mean of 42.3% and 48.8% respectively at an E: T ratio of 1:1 and by of 69.7% and 77.2%, at a ratio of 5:1. No reduction of p24+ cells was observed with the irrelevant TCR transduced cells at 1:1 and 23.5% at 5:1, whilst a 99.9% reduction was seen with KK10 TCR transduced cells at both ratios (Figure 4D). However KK10 TCR transduced cells showed no inhibition on HIV-1 infected CD4 T cells that were HLA B*2705 negative.

The virus inhibition experiments confirm that recognition of the RL9 peptide and HLA-E is TCR mediated and that HIV-1 infected cells present the RL9-HIV-1 peptide-HLA-E complex. The latter point is crucial to establish because priming of MHC-E restricted T cells appears to be more likely when the classical class I antigen processing pathway is modulated or bypassed *(7, 24).* HIV and SIV might do this to some degree enabling small amounts of HLA-E RL9 peptide reach the surface of infected CD4 T cells, sufficient to render them targets for T cell recognition and virus inhibition but insufficient to prime T cells *in vivo*.

The rhesus cytomegalovirus strain 68-1 (RhCMV68-1) vectored SIV vaccine described by Hansen et al *(9–11)* is the only current vaccine approach capable of arresting early SIV infection. Although not all animals are protected by this vaccine, the 57% level of protection is superior to all other experimental SIV or HIV-1 vaccines. Given the difficulty in producing a HIV-1 vaccine that stimulates broadly neutralizing antibodies *(25)* these results offer the hope of an alternative CD8+ T cell-based vaccine strategy, if findings from the RMs studies can be translated into humans. The key to the RhCMV68.1-SIV vaccine efficacy is the atypical CD8+ T cell response that it elicits *(7, 8)*, restricted by MHC class II or by MHC-E. This unusual T cell response is dependent on the lack of the orthologs of HCMV UL128/130 and UL146/147 in the viral vector *(7, 12)*. Recent data show that complete or partial restoration of these CMV genes in the RHCMV68-1 vector results in either a classical MHC-Ia restricted CD8 T cell response or a mixed MHC-Ia and MHC-II restricted T cell response and in both cases the protection is lost *(12)*. Furthermore, if the Rh67 gene, which delivers the VL9 signal peptide, is deleted from the RhCMV68-1 vaccine, the MHC-E restricted T cell response is lost concomitant with lack of protection against SIV challenge *(13)*. Therefore the MHC-E restricted T cells are critical to the effective anti-SIV T cell response.

We demonstrate here that it is possible to prime HLA-E restricted T cell response to an HIV-1 epitope in uninfected, HIV-1 negative humans. This is encouraging in the quest for an HIV-1 vaccine because it suggests the likelihood of being able to elicit T cell responses in humans, similar to the protective RM responses, using vaccines. The reduction of virus replication by HLA-E restricted TCR transduced into third party CD8+ T cells also offers a novel therapeutic approach to HIV-1 eradication.

## ACKNOWLEDGMENTS

We thank Professor Masafumi Takaguchi (University of Kumomotao) and Drs Dris Elatmioui and Thorbald van Hall for generous gifts of reagents. We dedicate this paper to the memory of our friend and co-author Enzo Cerundolo who sadly died in January 2020

## AUTHOR CONTRIBUTIONS

HY performed and designed the experiments, analyzed the data and contributed to writing of the manuscript. HY, HS and EB cloned the T cells. MR and PK and VC performed TCR sequencing and data analysis. MR made TCR J8 Jurkat transductants and primary CD8+ T cell transductants. SB made HLA-E-RL9 SCT. GG made HLA-E RL9 tetramers. ML, WM and XX made K562 HLA-E and TCR transductants. EJ and SD made tech TCR negative, CD8 positive, CD4 negative J8 Jurkat cell lines. SA and JBS made KK10 TCR plasmid construct. KF and LJP advised and helped design the studies. AJM directed the project and wrote the manuscript. AJM, GG and PB supervised the project. All authors read and approved the final version of the manuscript.

## DATA AND MATERIALS AVAILABILITY

All data associated with this study are present in the paper or Supplementary Materials.

## SUPPLEMENTARY MATERIALS LIST

### MATERIALS AND METHODS

#### Peptides

Synthetic 9 amino acid RL9HIV (RMYSPTSIL) and 11 amino acids RL9HIV-Gly-Cys (RMYSPTSILGC) peptides were generated by Fmoc (9-fluorenylmethoxy carbonyl) chemistry to a purity of 85% (Genscript, Hong Kong). All peptides were provided as lyophilized power. Following reconstitution to a final concentration of 200mM in DMSO, peptide stocks were aliquoted and stored at −80°C until required. A UV photolabile HLA-B leader-sequence peptide (VMAPRTLVL) incorporating a UV-sensitive 3-amino-3-(2-nitrophenyl)-propionic acid residue (J residue) substitution at the peptide p5 Arg residue was synthesized by Dris Elatmioui at LUMC, The Netherlands as previously described *(15).* This peptide, known herein as the 7MT2 peptide, was stored as lyophilized power at −80°C and reconstituted as required.

#### PCR-based site directed mutagenesis of the HLA-E*01:03 heavy chain

Position 84 Tyr to Cys mutagenesis of the HLA-E*01:03 heavy chain was performed by QuikChange II XL Site-Directed Mutagenesis Kit (Agilent, USA) using the following primers: Fw: 5’-CGGACGCTGCGCGGCTGCTACAATCAGAGCGAG-3’ and Rv: 5’-CTCGCTCTGATTGTAGCAGCCGCGCAGCGTCCG-3’]. A prokaryotic PET22b+ expression vector encoding HLA-E*01:03 heavy chain (residues 1-276) linked to a 15 amino acid biotinylation AviTAG was used as PCR template. Following mutagenesis and transformation into XL10 Gold bacteria, individual colonies were grown overnight in low salt Luria-Bertani (LB) broth containing 100μg/mL Carbenicillin. All plasmids were extracted with Spin miniprep kit (Qiagen, UK), and DNA Sanger sequencing confirmed their sequences.

#### Protein production

Both canonical and Tyr84Cys mutated HLA-E*01:03 heavy chains were expressed in *E. coli* BL21 (DE3) pLysS competent bacterial cells (Promega, UK). Single colonies inoculated in 1 Liter low salt Luria-Bertani (LB) broth containing 100μg/mL Carbenicillin were incubated overnight at 37°C. The following day, approximately 75mL of overnight starting cultures were transferred to fresh 3 x 1 liter Carbencillin-spiked LB broth. Following incubation to an OD600 of 0.5, protein expression was induced by the addition of 0.5mM IPTG. The cultures were subsequently incubated for a further 4 hours prior to bacterial pellet recovery via centrifugation at 1000g for 20 minutes at 4°C. Inclusion body proteins were extracted from bacterial pellets by sonication and homogenisation in a Triton-based buffer (Triton X-100, 50 mM Tris, 100mM NaCl, 0.1% Sodium Azide, 1 mM EDTA, 1mM DTT) followed by resuspension in a Tris-NaCl buffer (50mM Tris, 100mM NaCl, 1 mM EDTA, 1mM DTT). HLA-E*01:03 heavy chains were solubilized in 8M urea containing 50mM MES pH 6.5, 0.1mM EDTA, 0.1mM DTT and subsequently aliquoted at 10mg/mL and stored at −80°C until required.

#### Protein refolding and purification

Conventional and Tyr84Cys mutated HLA-E*01:03 heavy chains were refolded using standard MHC refolding methods *(1, 26)* but with slight modifications. In brief, β2m (2μM final concentration) was refolded in MHC Refold Buffer (100mM Tris pH8.0, 400mM L-Arginine monohydrochloride, 2mM Ethylenediamineteraacetic acid, 5mM reduced Glutathione and 0.5mM oxidised Glutathione) for 30 minutes at 4°C, following which either 30μM RL9HIV-GC peptide (for Cys trapped refolds) or 50μM RL9HIV peptide (for conventional refolds) was added. Conventional or Tyr84Cys mutated HLA-E*0103 heavy chains were subsequently pulsed into the refolding buffers respectively, up to a final concentration of 1μM. After 72 hours, all refolds were filtered using 22μm cellular nitrate membrane (GE Healthcare, UK) before concentration using Vivaflow 50 (Sartorius, Germany) and Ultra-15 10-kDa cut-off centrifugal units (Sartorius, UK). The samples were subsequently buffered exchanged, using Sephadex G-25 PD10 columns (GE Healthcare, UK) into 10mM Tris for overnight AviTAG biotinylation using the BirA enzyme (Avidity, USA) according to the manufacturer’s instructions. Correctly refolded complexes were purified by size exclusion fast protein liquid chromatography (FPLC) into 20mM Tris pH8 and 100mM NaCl buffer using a HiLoad 16/600 Superdex 75pg column. Correctly folded β2m-HLA-E*01:03-peptide complexes were retrieved, concentrated to 2mg/mL and snap frozen for subsequent tetramer generation.

#### Protein thermal melt analysis

The thermal stability of refolded peptide/HLA-E complexes was evaluated by heat-induced fluorescent dye incorporation using the Protein Thermal Shift™ Dye kit (Applied Biosystem, USA). Protein Thermal Shift ROX Dye and Protein Thermal Shift Buffers were freshly prepared and aliquoted into MicroAmp Fast Optical 96-well plates (Applied Biosystem, China), according to the manufacturer’s instructions. 5μg of refolded HLA-E material was added to individual wells, and buffer control wells lacking protein served to monitor background fluorescent signals. Samples and controls were set up in duplicate, and at least 3 biological runs were evaluated per test sample. All assays were performed on an Applied Biosystem Real-Time 7500 Fast PCR System, with a temperature ramp from 25 to 95°C and 1°C intervals. Single thermal melt reads of inflection point data were determined using Protein Thermal Shift Software v1.3. Means and standard deviations of the means are reported.

#### Generation of UV exchange RL9-loaded HLA-E

Refolding of the VL9-based UV sensitive (7MT2) peptide with HLA-E and β2m was performed as previously described *(15, 27).* Protein concentration and biotinylation was carried out as per the method described in **Protein refolding and purification**. For UV-mediated peptide exchange reactions, 10 wells comprising 0.5μM (~25mg/mL) of HLA-E-7MT2 monomer were incubated with 150μM RL9 peptide in polypropylene V-shaped 96-well plates (Greiner Bio-One, Austria). UV exchange buffer (20mM Tris, pH 7.4, 150mM NaCl) was added to each well to adjust the final reaction volumes to 125μL. UV exchange samples were incubated under a Camag UV cabinet with a long-wave 366nm UV lamp for 60 minutes on ice. Following photo-illumination, the samples were centrifuged at 4000g for 20 minutes to remove aggregated material. Aggregate-cleared samples were pooled and conjugated to fluorescent dyes as described below (**Tetramer generation and staining protocol**)

#### Tetramer generation protocol

Disulfide trapped and UV-peptide exchange HLA-E*01:03-RL9 tetramers were generated via conjugation to streptavidin-bound APC (Biolegend, San Diego) or BV421 (Biolegend, San Diego) at a Molar ratio of 4:1 as previously described *(1)*.

#### Cell lines and primary cells

The MHC-I null cell line K562 transfected with HLA-E*01:03 (K562-E line) was generously provided by Thorbald van Hall (Leiden University Medical Centre) *(17).* The 721.221 HLA-class I deficient cell line transfected with CD4 (721.221-CD4) was generously provided by Masafumi Takiguchi, University of Kumomoto, Japan *(28, 29).* PBMCs were isolated from HIV negative donor leukapheresis cones (NHS Blood and Transplant, UK) by density gradient separation. CD4+ and CD8+ T-cells were enriched from PBMC by positive selection using magnetic bead according to the manufacture’s instructions (MACS, Miltenyi Biotech, Surrey, UK).

#### Transduction of K562 cells with HLA-E-RL9 single chain trimers (SCT) construct (K562-E-RL9) and disulfide trapped (K562-D-RL9) constructs

Single chain trimers (SCT) of HLA-E*01:03 with the RL9HIV peptide (RMYSPTSIL) constructs were generated as previously described *(1, 27).* A disulfide “trap” was engineered into the SCT by mutating position 84 of HLA-E to cysteine and changing the sequence of the first flexible linker (between the peptide and beta2-microglobulin) to GCGGSGGGGSGGGGS *(20).* The constructs were ligated into the retroviral vector, pMSCV-GFP (Addgene). Retroviral particles were produced by mixing 2μg of the HLA-E plasmids with 0.5μg of pCMV-VSV-G (Cell Biolabs) and 200μl of OPTI-MEM (Gibco) for 5mins at room temperature. 7μl of X-tremeGENE HP Transfection reagent (Roche) was added and incubated at 37°C, 5% CO_2_ for 15mins. This transfection solution was added to PlatGP cells (Cell Biolabs) and incubated overnight at 37°C, 5% CO_2_. Retroviral particles were harvested after 24 hours and stored for 3 days after initial transfection. Twenty-four well plates pre-coated with 15μg/ml RetroNectin were blocked with 2% BSA, PBS. 1 x 10^6^ K562 cells were transduced in each well with 2 ml of retrovirus supernatant by centrifugation at 100g, for 2 hours at 32°C. HLA-E transduced K562 cells were further purified by cell sorting, based on the expression of HLA-E as determined by staining with the 3D12 mAb clone (BioLegend).

#### Tetramer staining protocol

Cells were stained with disulfide trapped HLA-E-RL9 tetramer or UV exchanged RL9 tetramer both conjugated to APC, at 0.5ug per 1×10^6^cells in 100μl MAC buffer (PBS with 2mM EDTA and 0.5% BSA) at room temperature (RT) for 45 minutes in the dark. After washing with PBS, cells were further stained with CD8-BV421 antibody (BioLegend) and Live/Dead Fixable Aqua dye (Thermo Fisher Scientific) in 100 μl PBS for 30 min at RT in the dark. After another PBS wash and fixation with 2% paraformaldehyde, cells were acquired using an LSR Fortessa (BD Biosciences) and the data analyzed using FlowJo software v10.3 (Tree Star).

#### In vitro priming of HLA-E restricted RL9HIV specific CD8+ T cells

On day 0, 100 to 150×10^6^ freshly isolated PBMCs were plated at 10^7^/ml in 6-well plates in AIM-V medium (Invitrogen) with a dendritic cell (DC) differentiation cytokine cocktail of GM-CSF (1000U/ml, Miltenyi Biotech Ltd) and IL-4 (500U/ml, Miltenyi Biotech Ltd). On day 1, DC maturation stimuli of TNF-α (1000U/ml, R&D Systems), IL-1β (10ng/ml, R&D Systems) and prostaglandin E_2_ (PGE_2_ 1μM, Merck) were added together with RL9HIV peptide (20μM, GenScript), IL-7 (5ng/ml, R&D Systems) and IL-15 (5ng/ml, R&D Systems). On day 6, IL-2 was added at a concentration of 500IU/ml. HLA-E RL9 tetramer staining was evaluated on day 9. In selected experiments, cells were further stimulated with irradiated (120 Gy) K562-D-RL9 cells for 7 days.

#### Cloning of HLA-E restricted RL9HIV specific CD8+ T cells

After RL9HIV priming, PBMCs were stained with an APC conjugated disulfide trapped HLA-E RL9 tetramer at 5ug per 5×10^7^ cells in 500μl MAC buffer at RT for 45 minutes in the dark. After a PBS wash, cells were further stained with anti-CD3-APC-Cy7, anti-CD4-PerCP-Cy5.5, anti-CD8-BV421, anti-CD94-FITC (All BioLegend) and the dump markers Live/Dead Fixable Aqua, anti-CD56-BV510 (BD Biosciences) for 30 min at RT in the dark. Tetramer+/CD3+/CD8+/CD4-/CD56-/CD94-/live subsets were sorted using a FACS Aria III (BD Biosciences). Sorted tetramer+ cells were seeded at 0.4 cells/well into 384-well plates (Corning) with phytohemagglutinin (PHA 1mg/mL, Remel) and irradiated (45 Gy) allogeneic feeder cells from 3 different HIV-negative donor leukapheresis cones (10^6^ feeder cells/mL) in RPMI 1640 glutamine [-] medium (Invitrogen) supplemented with non-essential amino acids (1%, Invitrogen), sodium pyruvate (1%, Invitrogen), glutamine (1%, Invitrogen), b-mercaptoethanol (0.1%, Invitrogen), penicillin/streptomycin (1%, Invitrogen) (RPMI 1640 complete media (RPMI 1640 CM)) with pooled AB human sera (10%, UK National Blood Service) and IL-2 (500 IU/mL). After 10 days, T cell clones were visually identified and transferred into 96-well round-bottom plates (Corning). An aliquot of each clone was stained with HLA-E-RL9 disulfide trapped tetramer and anti-CD3-APC-Cy7, anti-CD8-BV421, anti-CD4-PerCP-Cy5.5 anti-CD94-FITC antibodies and dump markers Live/Dead Fixable Aqua, anti-CD56-BV510 to confirm RL9HIV specificity.

#### TCR sequencing

RNA was extracted from the T cell clones using a RNeasy Micro Kit (Qiagen), following manufacturer’s instructions. TCR cDNA was generated by template-switch reverse transcription, using a template switch oligo, and primers specific to the constant regions of Trac (5’-TCAGCTGGACCACAGCCGCAG-3’) and Trbc (5’-CAGTATCTGGAGTCATTGA-3’) genes, and SMARTScribe Reverse Transcriptase (Takara). Two subsequent rounds of nested PCR using Phusion High-Fidelity PCR Master Mix (NEB) amplified TCR DNA. One last PCR was performed to add the Illumina adaptors and indexes. TCR libraries were sequenced using an Illumina Miseq Reagent Kit V2 300-cycle on the Illumina Miseq platform. FASTQ files were demultiplexed and TCR sequences analyzed using MiXCR software *(30).* Post analysis was performed using VDJtools *(31)*.

#### Generation of the J8 CD8^+^ Jurkat T-cell line using CRISPR-Cas9

For ablating Jurkat TCRα (TRAV8-4*01), TCRβ (TRBV12-3*01) and CD4 (UniProtKB P01730) gene expression using CRISPR-Cas9, guides were designed and selected for high specificity with minimal off target activity using Benchling (hg38 reference genome) and CRISPOR (Table S2). For each gRNA sequence, complemenatry oligonucleotides with appropriate overhangs were annealed, and ligated into the LentiCRISPRv2 plasmid (Addgene plasmid 52961) using the dual BsmBI restriction sites, as described elsewhere *(32, 33)*.. For stable insertion of CD8α (aa 22-235, UniProtKB P01732) and CD8β (aa 22-210, UniProtKB P10966), the genes were cloned into a pHR plasmid backbone modified with the RPTPμ phosphatase signal peptide (UniProtKB P28827). Lentiviral particles were produced by lipid nanoparticle transfection into HEK293T cells (GeneJuice, Novagen). Briefly, 0.5 μg of pHR or LentiCRISPRv2 plasmids were co-transfected with 0.5 μg of envelope (pMD2. G) and 0.5 μg packaging (p8.91) plasmids. Medium containing viral particles from the HEK293T cells was 0.22 μm filtered two days later, pooled when necessary, and added directly to 1×10^6^ Jurkat cells. Three to seven days later, the cells were sorted by FACS and/or treated with puromycin at a concentration of 1 μg/ml in complete medium for 3 days, before moving to 10 μg/ml for 3 additional days (**Figure S4**).

#### TCR transduction into J8 Jurkat cells and primary CD8+ T cells

TCR alpha and beta VDJ regions were amplified by PCR from the DNA generated during the preparation of TCR sequencing libraries. These products were assembled into a pHR-SIN backbone with the murine TCR alpha and beta constant regions to avoid formation of hybrid TCRs with endogenous TCRs and for ease of detection, using the HiFi DNA Assembly cloning kit (NEB). Sanger sequencing confirmed correct plasmid sequences. Lentiviruses were produced by transfecting the TCR-containing plasmid plus pMDG-VSVG, and pCMV-dR8.91 packaging plasmids into HEK 293T cells, using the transfection reagent TurboFectin (Origene). Lentiviral supernatants were collected 48h after transfection, centrifuged at 2000rpm to remove cellular debris and transferred to Retronectin (Takara Bio) treated 48-well plates. The plates were centrifuged 1.5h, at 2000xg to facilitate virus binding and supernatant was removed. Primary T cells were isolated from PBMC by positive selection using MACS beads (Miltenyi) and activated for 2 days with 1:1 of CD3/CD28 Dynabeads (Thermo Fisher) in RPMI medium supplemented with 1% non-essential amino acids, 1% sodium pyruvate, 1% glutamine, 1%HEPES, 1% pen-strep 0.1%β-mercaptoethanol (Invitrogen), 5% pooled AB human sera (UK National Blood Service), 500U/mL IL-2 (University of Oxford), and 10ng/mL rhIL-5 (Peprotech). Activated T cells were transferred to lentivirus-coated plates at 0.25×10^6^cells/mL and cultured for 4 days. Mouse TCRβ+ CD8+ cells were purified by flow cytometry (BD Fusion) and expanded for a further 17 days before usage in subsequent assays. J8 CD8+ Jurkat T-cells were further modified to express a NFAT-eGFP reporter system using the pSIRV-NFAT-eGFP *(34).* The reporter J8 Jurkat cells were transduced with TCRs by adding 500μL of lentiviral supernatant to 500 μL of cell suspension at 1×10^6^cells/mL in 6-well plates.

#### Construction of HLA-B27:05-restricted HIV Gag_263-272_ (KK10) TCR CD8+ T cell transductants

CD8+ T cell transductants targeting the HIV-1 Gag_263-272_ KK10 epitope (KRWIILGLNK) restricted by HLA-B*27:05 were based on the published C12C clone *(35)*. To construct this full-length TCR construct, we utilized the published C12C CDR3α and CDR3β sequences and combined these with nucleotide sequence provided by IMGT for TRBV6-5, TRBJ1-1, TRAV14, and TRAJ21. After murinization and cysteine modification of the constant domains of the TCR and insertion of an additional cysteine bridge, the complete sequence of both TCR chains was constructed with a 2A sequence for bi-cistronic expression (Genscript, Piscataway Township, NJ, USA) and cloned into the pMP71 backbone. To produce retroviral supernatants, the TCR construct was transfected into the embryonal kidney cell line 293Vec-RD114 (BioVec Pharma, Québec, Canada). Collected supernatants were then purified via centrifugation on a 20% sucrose gradient. Viral transduction of activated CD8+ T cells was done by magnetofection using Viromag Viral Transduction reagent (Oz Biosciences, Marseilles, France) according to manufacturer’s protocol. CD8+ T cell transductants were then cultured in X-vivo 15 (Lonza, Basel, Switzerland) supplemented with 10% FBS and 200U/ml IL-2 until use in assays.

#### Evaluation of IFN-γ, TNF-α, CD107a/b and CD137 upregulation

Clone cells were washed and left in fresh RPMI 1640 CM (5% AB serum) without IL-2 to rest for 5 hours or overnight before being stimulated with RL9 peptide pulsed K562-E (50μM, 20-24 hours at 27°C), K562-E-RL9 or K562-D-RL9 cells at a clone: K562 cell ratio of 1:3 for 1 hour, followed by addition of 5μg/ml Brefeldin A (Biolegend) and 5μg/ml GolgiStop (BD Biosciences) for an additional 8 hours at 37°C. For CD107 staining, anti-CD107a-BV421 and anti-CD107b-BV421 (Biolegend) antibodies were added at the beginning of the co-culture. After 9 hours incubation, cells were washed with PBS and stained with Live/Dead Fixable Aqua, anti-CD8-PerCP-Cy5.5 and anti-CD3-APC-Cy7 for 30 min at RT first, then fixed/permeabilized with Cytofix/Cytoperm 1x Solution (BD Biosciences) for 10 min at 4°C, and stained in Permwash 1x Solution (BD Biosciences) with anti-TNFα-PE, anti-IFN-γ-FITC and anti-CD137-BV650 (All BioLegend) for 30 min at RT. After being washed with PBS and fixed with 2% paraformaldehyde, samples were acquired using an LSR Fortessa (BD Biosciences) and analyzed using FlowJo software v10.3 (Tree Star). In the selected assays to determine HLA-E restriction, K562-E cells were pre-incubated with the VL9 canonical signal peptide (VMAPRTLVL, 50μM, 3 hours at 27°C) prior to addition of RL9 peptide.

#### Activation of RL9TCR transductants

Jurkat J8 cells transduced with RL9 TCR were labelled with CellTrace Violet Dye in accordance with the manufacturer’s instructions (ThermoFisher Scientific) for easy identification during analysis. They were then stimulated with RL9 peptide pulsed K562-E (50μM, 20-24 hours at 27°C), K562-E-RL9 or K562-D-RL9 cells at clone: K562-E cell ratio of 1:3 for 8 hours at 37°C. Cells were washed with PBS and stained with Live/Dead Fixable Far Red stain (ThermoFisher Scientific), anti-CD8-PerCP-Cy5.5 and anti-CD69-PE for 30 min at RT. Cells were washed with the PBS and then fixed with 2% paraformaldehyde before flow cytometry analysis. Primary CD8+ transductants were washed and rested in RPMI 1640 CM media with 10% human serum for minimal 5 hours or overnight prior stimulated with RL9HIV peptide pulsed autologous B cells (50μM, 2 hours at 37°C) at a transductant: B cell ratio of 1:2 for intracellular TNFα cytokine and CD137 staining as described earlier.

#### Viral inhibition / infected cell elimination assay (VIA)

PBMCs were stimulated with anti-human CD3 at 100ng/ml (clone OKT3, TONBO Biosciences) in RPMI 1640 CM supplemented with 5% AB human serum and IL-2 (100 IU/ml) for 5 days. CD4+ cells were enriched from activated PBMC by positive selection using anti-CD4 magnetic beads according to the manufacturer’s instructions (MACS, Miltenyi Biotech, Surrey, UK). Activated CD4+ cells or 721.221-CD4 cells were infected with the HIV-1 NL4.3 virus obtained from the Programme EVA Centre for AIDS Reagents (National Institute for Biological Standards and Control (NIBSC), a centre of the Health Protection Agency, UK.) at a multiplicity of infection of 1 × 10^-2^ by spinoculation for 2 hours at 27°C, as described previously *(36).* HIV-1 NL4.3-infected target cells (primary CD4+ T-cells or 721.221-CD4 cells) were washed with RPMI 1640 CM and cultured in triplicate (1 × 10^5^ cells/well) in RPMI 1640 CM supplemented with 5% AB serum and IL-2 (50 IU/ml), either alone or with RL9 clone cells or primary CD8+ transductants for 5 days at various Effector: Target (E: T) ratios. An EBV clone (B*0801 restricted RAKFKQLL specific) or non-transduced primary CD8+ T cells were used as a control for RL9 specificity. After 5 days of coculture, cells were collected and stained with Live/Dead Fixable Aqua before permeabilizing with BD fix/perm solution for intracellular HIV Gag p24 (Beckman Coulter, UK) staining followed by staining with anti-CD3-APC-Cy7, anti-CD8-BV421, anti-CD4-PerCP-Cy5.5 antibodies. The frequency of infected cells was determined by intracellular staining for Gag p24 Ag, optimized for sensitivity and specificity, as described previously *(21, 36).* To demonstrate the presentation of RL9 epitopes by HLA-E on HIV-1 NL4.3 infected CD4+ T cells, selected experiments were conducted with the addition of excess competing VL9 canonical signal peptide (50μM) in the coculture of targets and effectors.

Viral inhibition / infected cell elimination was calculated by normalising to data obtained with no effectors using the formula: (fraction of Gag+ cells in CD4+ T-cells cultured alone - fraction of Gag+ in CD4+ T-cells cultured with CD8+ clone cells) / fraction of p24+ cells in CD4+ T-cells cultured alone × 100%. In selected experiments, CD8+ clone T cells were analyzed for expression of the activation marker CD137 using BV421-conjugated antibodies (BD Biosciences) at 24 hours post effector and target co-culture.

The VIA was set up with a minimum of 3 replicates for each culture condition. Cells from each culture condition were harvested and pooled for intracellular p24 staining to reach the required acquisition of at least 10000 viable target cells for each target and effector coculture.

#### Statistical analysis

Statistical analysis was performed using GraphPad Prism software (version 6.0 or later). Data with skewed distributions were analyzed with the non-parametric test, Wilcoxon signed rank test. Where a normal distribution was observed, data were analyzed with parametric tests (Repeated Measures 2-way ANOVA with Tukey’s multiple comparisons tests).

## Supplemental Figures

**Figure S1.**
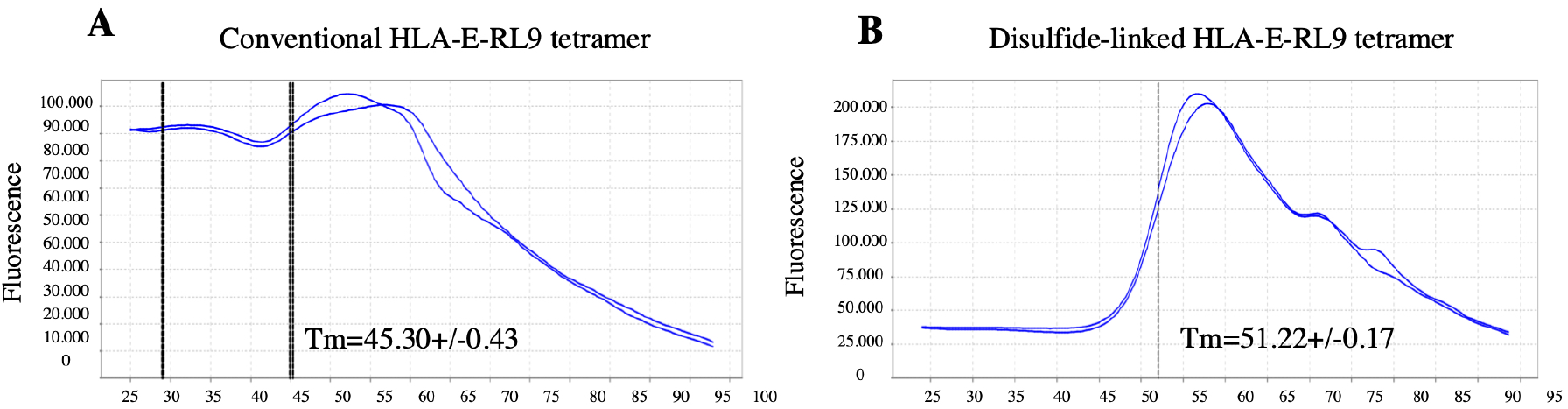
Enhanced stability of disulfide-linked RL9 HLA-E complexes versus conventionally refolded RL9-HLA-E material, illustrated by thermal melt analysis. Thermal melt analysis of (**A**) conventional and (**B**) disulfide-linked RL9-HLA-E complexes as measured by fluorescent (ROX) dye incorporation using the Applied Biosystems Thermal Shift Assay. The raw melt curve data is illustrated, with the Y-axis denoting fluorescence intensity (arbitrary units) and the X-axis depicting time (minutes). The inflection point of each melt curve, defined as the derivative melting temperature (TmD), was calculated automatically using Protein Thermal Shift Software v1.3. Two technical replicates were tested per run, and two biological replicates were performed for each tetramer. Representative data from a single assay illustrating Tm Derivative (TmD) values (denoted as black vertical lines on curves) +/− standard deviations are reported.

**Figure S2.**
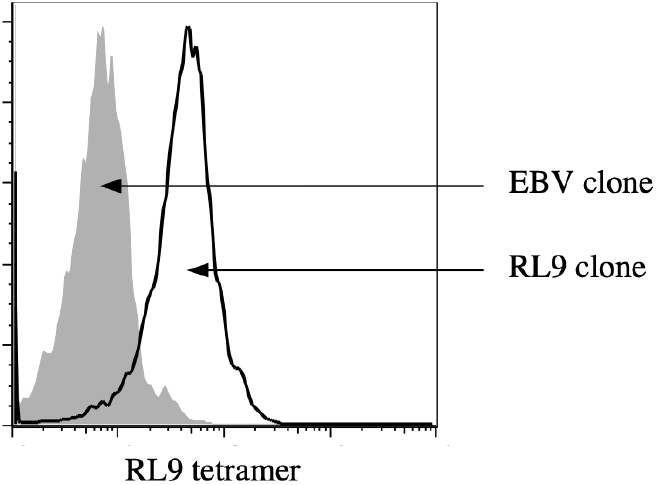
Representative FACs plot of a RL9 clone stained with HLA-E-RL9 disulfide-linked tetramer, fixed without wash. 1 million RL9 clone cells were stained with 0.2μg HLA-E-RL9 disulfide-linked tetramer for 40 minutes at RT prior to fixation with 2% paraformaldehyde for 15 minutes in the absence of washing step before acquisition on a LSR Fortessa. The tetramer failed to bind a B*08:01 restricted EBV-specific CD8+ T cell control clone generated from the same donor.

**Figure S3.**
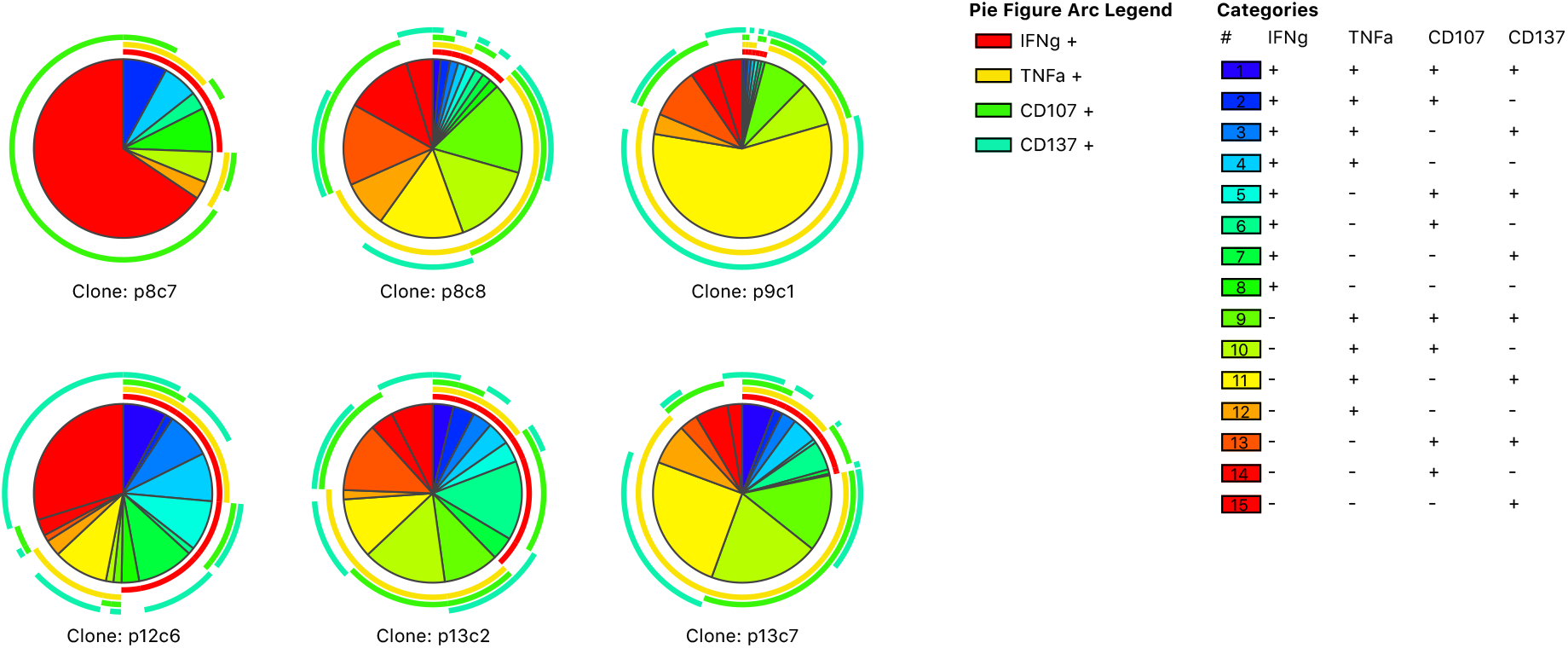
Multiple function analysis of HLA-E restricted RL9HIV-specific CD8^+^ T cell clones. RL9HIV specific responses are divided into 15 distinct subpopulations with any combination of IFNγ, TNFαsecretion and CD107a/b, CD137 expression using SPICE analysis. Pie charts represent combination of responses of 6 clones. The arcs highlight one single response among the total responses.

**Figure S4.**
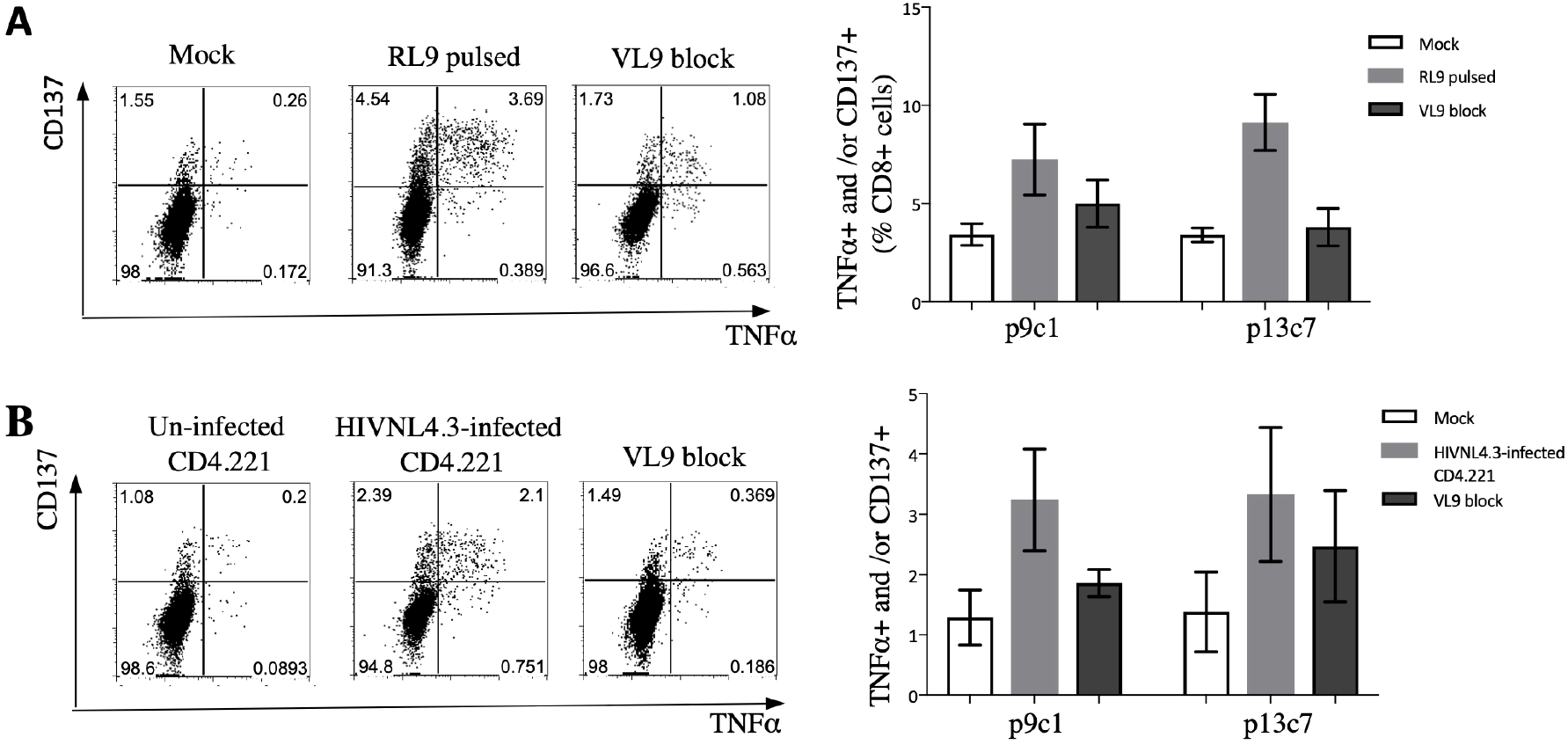
Primary CD8+ T cells transduced with RL9 TCRs are activated by RL9 stimulation and can reduce the proportion of HIV-infected CD4+ T cells. (**A**) CD8+ transductants were activated by exposure to autologous B cells pulsed with RL9 peptide or (**B**) HIVNL4.3-infected CD4.221 cells, indicated by TNFα secretion and up-regulation of CD137. Responses were partially blocked by competitive inhibition with the signal peptide VL9. Gag p24+ cells were gated on CD3+CD8-T cells. Horizontal lines indicated means. Error bars indicated SD. Data shown is representative of three independent experiments.

**Table S1.**
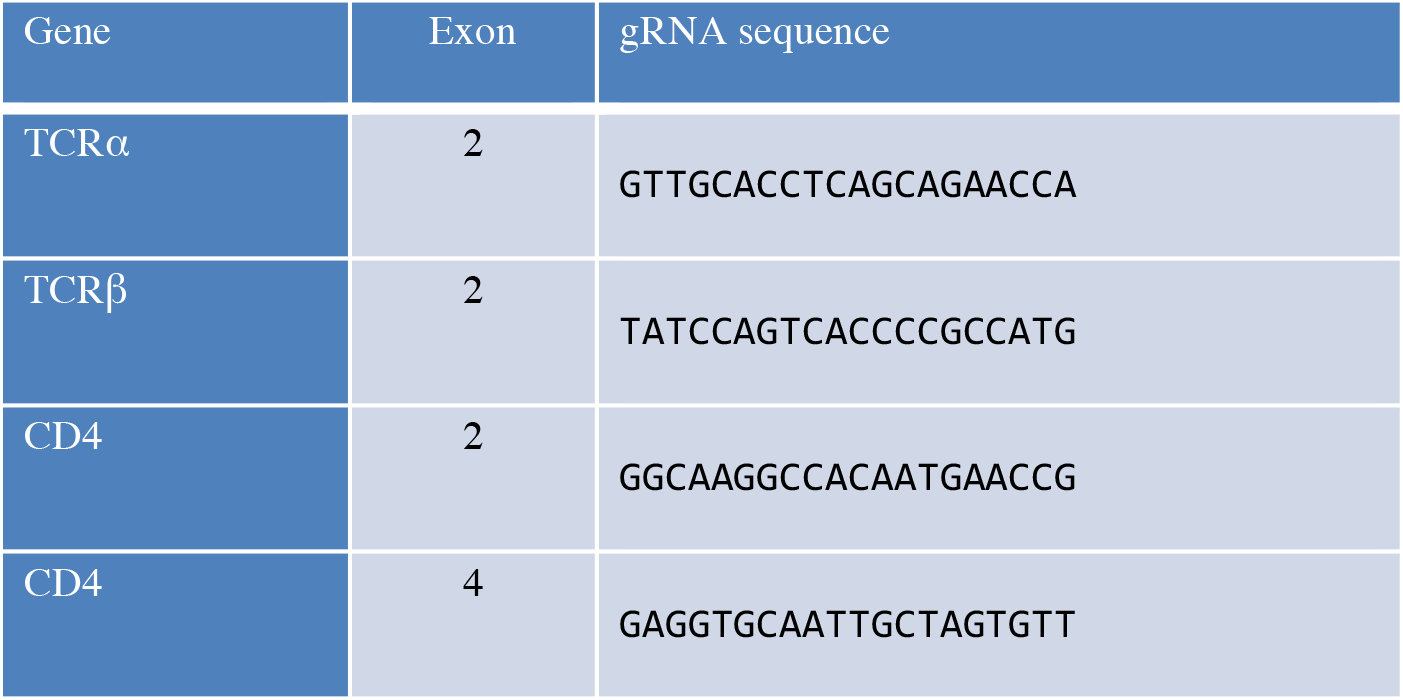
CRISPR guides for TCR and CD4 deletion in Jurkat cells.

**Figure S5.**
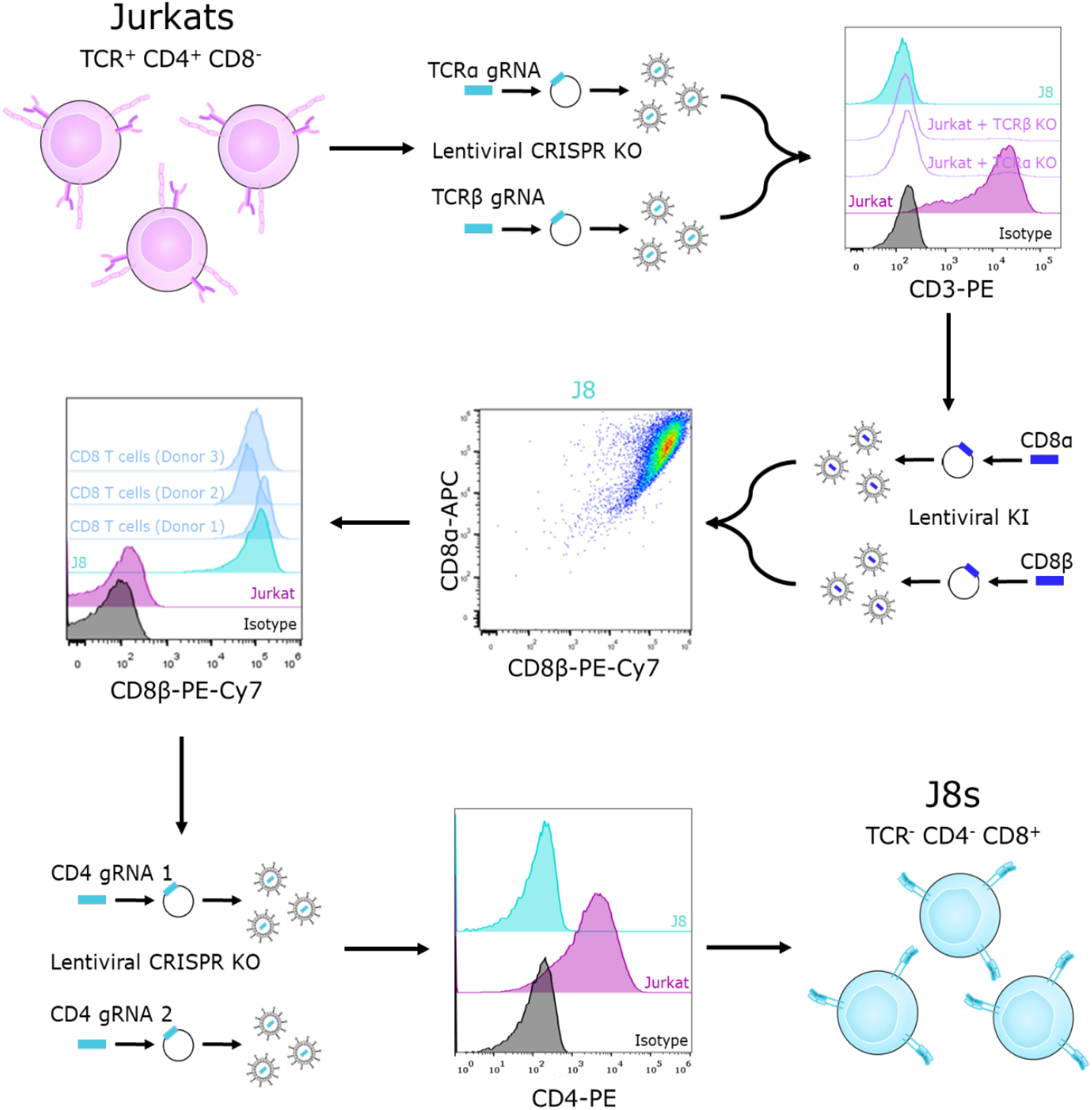
Creation of the CD8^+^ Jurkat (J8) T cell line. Jurkat (leukemic CD4^+^) T-cells were transduced simultaneously with lentiviruses expressing Cas9 and gRNA specific for the endogenous TCRα and TCRβ genes, using LentiCRISPRv2. Simultaneous knockout was possible owing to the efficiency of the individual guides (>90%). Cells were then sorted in bulk to remove residual CD3^+^ cells, with stable insertion of both guides further selected for using puromycin selection for one week. The CD8α/β chains were then introduced into the cell-line and CD8α/β expression matched by cell sorting to human-PBMC derived CD8^+^ T cell levels. Lastly, human CD4 expression was ablated using two gRNA-Cas9 containing lentiviruses, with cells sorted in bulk on negative CD4 expression without the need for further puromycin treatment, giving the J8 cell-line.

